# LncRNA GAS5 attenuates fibroblast activation through inhibiting Smad3 signaling

**DOI:** 10.1101/2020.02.01.930412

**Authors:** Rui Tang, Yung-Chun Wang, Xiaohan Mei, Ning Shi, Chengming Sun, Ran Ran, Gui Zhang, Wenjing Li, Kevin F. Staveley-O’Carroll, Guangfu Li, Shi-You Chen

## Abstract

Transforming Growth Factor β (TGF-β)-induced fibroblast activation is a key pathological event during tissue fibrosis. Long noncoding RNA (lncRNA) is a class of versatile gene regulators participating in various cellular and molecular processes. However, the function of lncRNA in fibroblast activation is still poorly understood. In this study, we identified growth arrest-specific transcript 5 (GAS5) as a novel regulator for TGF-β-induced fibroblast activation. GAS5 expression was downregulated in cultured fibroblasts by TGF-β and in resident fibroblasts from bleomycin-treated skin tissues. Overexpression of GAS5 suppressed TGF-β-induced fibroblast to myofibroblast differentiation. Mechanistically, GAS5 directly bound Smad3 and promoted Smad3 binding to PPM1A, a Smad3 dephosphatase, and thus accelerated Smad3 dephosphorylation in TGF-β-treated fibroblasts. In addition, GAS5 inhibited fibroblast proliferation. Importantly, local delivery of GAS5 via adenoviral vector suppressed bleomycin-induced skin fibrosis in mice. Collectively, our data revealed that GAS5 suppresses fibroblast activation and fibrogenesis through inhibiting TGF-β/Smad3 signaling, which provides a rationale for an lncRNA-based therapy to treat fibrotic diseases.

## Introduction

Tissue fibrosis is a wound healing progress following injury, during which normal parenchymal tissue is gradually replaced by connective tissue, accompanied by deposition of extracellular matrix (ECM) components (7, 51, 53). The pathogenesis of fibrosis is initiated in the residential area where local injury triggers innate immune response and causes recruitment of immune cells (26, 32, 40). Pro-fibrotic cytokines or growth factors such as transforming growth factor β (TGF-β) secreted from circulating monocytes and residential macrophages induce fibroblast activation through fibroblast-myofibroblast transition (6, 22, 37, 38). Activated fibroblasts proliferate and thus increase tissue mass. Meanwhile, myofibroblasts express extracellular matrix proteins such as collagen 1A (Col1A), the main component of excessive ECM deposition in human fibrotic diseases such as skin, cardiovascular, pulmonary, and liver fibrosis. (57, 58).

TGF-β signaling regulates a wide spectrum of cellular processes, including cell differentiation, proliferation, migration, and apoptosis (33, 35). TGF-β induces fibrosis through inducing fibroblast trans-differentiation into collagen-producing myofibroblasts and increasing fibroblast proliferation (5, 29). TGF-β transduces its signal mainly through Smad-dependent pathways. TGF-β binds and activates TGF-β receptors (TβRs), leading to Smad phosphorylation and translocation into nuclei where it activates the transcription of target genes (14). However, how Smad signaling is precisely modulated during TGF-β-induced fibroblast activation and how TGF-β coordinates fibroblast transdifferentiation and proliferation during fibrogenesis are not completely understood.

Long non-coding RNAs (lncRNAs) are a class of versatile regulators which regulate gene expression at various levels ranging from chromatin modulation to protein degradation (25). Growth Arrest Specific 5 (GAS5) is a well-known lncRNA that suppresses cell proliferation and induces apoptosis (27, 36). Our previous studies have identified GAS5 as a regulator in both Smad3-dependent cell differentiation and p53-dependent cell survival (47, 49). Since TGF-β signaling regulates both Smad-dependent fibroblast-myofibroblast transition and fibroblast proliferation, we hypothesized that GAS5 may be involved in TGF-β-induced fibroblast activation and fibrogenesis.

In the current study, we firstly observed that GAS5 was downregulated in TGF-β-treated fibroblast cells and bleomycin-injected skin tissues. By both gain-of- and loss-of-function studies, we found that GAS5 suppressed TGF-β-induced fibroblast-myofibroblast transition through modulating Smad3 signaling. Mechanistically, GAS5 directly bound to Smad3, which enhanced phosphatase PPM1A binding to Smad3, and thus accelerated Smad3 dephosphorylation. GAS5 also inhibited fibroblast proliferation by blocking c-Jun N-terminal kinase (JNK) signaling pathway. In vivo, local delivery of adenoviral vector expressing GAS5 inhibited bleomycin-induced skin fibrosis in mice.

## Material and Methods

### Cells and Reagents

NIH-3T3 cells were purchased from American Type Culture Collection (ATCC). Cells were maintained at 37 °C in a humidified 5% CO2 incubator in Dulbecco’s Modified Eagle’s Medium (GIBCO, CA, USA) containing 10% fetal bovine serum (GIBCO, CA, USA), 100 units/ml penicillin and 100 μg/ml streptomycin. TGF-β1 was obtained from R&D Systems (Minneapolis, MN). mGAS5 siRNA (n251731) was purchased from Life Technologies (Gaithersburg, MD). Smad3 inhibitor SIS3 was purchased from Sigma-Aldrich (St. Louis, MO, USA). PPM1A (PA5–29275) antibody was purchased from ThermoFisher Scientific (Pittsburgh, PA)(15). Akt (4691S), phospho-Akt (9271S), JNK (9252S), phospho-JNK (9251S), p38 (9212S), phospho-p38 (4511S), Smad3 (9523S), phospho-Smad3 (9520S), and Smad4 (38454) antibodies were purchased from Cell Signaling (Danvers, MA, USA)(48, 49). GAPDH (G8795) and α-SMA (A2547)antibodies were purchased from Sigma-Aldrich (St. Louis, MO, USA)(43). PCNA (sc-56) and Type I Collagen (Col1A) (sc-25974) antibodies were purchased from Santa Cruz Biotechnology (Dallas, TX, USA) (43, 47). Smad3 overexpression plasmid was constructed by subcloning human Smad3 cDNA into pcDNA3.0 backbone plasmid (Addgene). GAS5 adenoviral vector was constructed by inserting mouse GAS5 cDNA into pShuttle-IRES-hrGFP-1 vector (Agilent), and adenovirus was packaged as described previously (49). All constructs were confirmed by Sanger sequencing.

### Primary mouse skin fibroblast preparation

Skin tissues of C57BL/6J mice (The Jackson Laboratory) were dissected, cut into small pieces, and digested in 5 ml tissue digest media (3.5 ml HBSS-Ca2+ free, 0.5 mL Trypsin-EDTA (0.25%), 5 mg Collagenase IV (Worthington), 25 U Dispase (Corning)) in a hybridization chamber with rotation at 37°C for 30 minutes. Digestion was then neutralized by adding 5 ml ice-cold Quench Solution (4.5 ml L15 media, 0.5 mL FBS, 94 µg DNase). Single cell suspensions were generated by filtering through a 40 uM cell strainer, spinning down at 500 rcf for 5 minutes followed by washing with PBS twice. Skin fibroblasts were resuspended and cultured in Dulbecco’s Modified Eagle’s Medium (GIBCO, CA, USA) containing 10% fetal bovine serum (GIBCO, CA, USA), 100 units/ml penicillin and 100 μg/ml streptomycin.

### Animals and skin fibrosis model

All animals were housed under conventional conditions in the animal care facility and received humane care in compliance with the Principles of Laboratory Animal Care formulated by the National Society for Medical Research and the Guide for the Care and Use of Laboratory Animals. Bleomycin-induced dermal sclerosis/fibrosis was generated following previously published protocol (43). 8-10 week old male C57BL/6 mice were injected subcutaneously with bleomycin (0.02U) in PBS every other day for 14 or 28 days. PBS was injected as control. Adenovirus (1×108 pfu per mouse) expressing GFP or mGAS5 was injected twice on the first day and 14 days following the first bleomycin injection. After 2 or 4 weeks, the animals were euthanized by CO2 asphyxiation and cervix dislocation. The skin areas with bleomycin injection were removed and processed for biochemical or histological analysis. All animal surgical procedures were approved by the Institutional Animal Care and Use Committee of the University of Georgia.

### Quantitative RT-qPCR (qPCR)

Total RNA was extracted from cells or tissues using Trizol reagent (Life Technologies, Gaithersburg, MD) and reverse-transcribed to cDNA using iScript™ cDNA Synthesis Kit (Bio-Rad, Hercules, CA). qPCR was performed using a Stratagene Mx3005 qPCR thermocycler (Agilent Technologies, La Jolla, CA). All reactions including no template controls were run in triplicates. After the reaction, the CT values were determined using fixed threshold settings. LncRNA expression was normalized to Cyclophilin (CYP). Primers used in this study were listed in Supplementary Table 1.

### RNA immunoprecipitation (RIP) assay

RIP assay was performed as described (49). Cells at 80-90% confluence in 15 cm2 culture dishes were fixed with 1% Paraformaldehyde (PFA) and lysed in FA lysis buffer (50 mM HEPES, 140 mM NaCl, 1 mM EDTA, 1% (v/v) Triton X-100, 0.1% (w/v) sodium deoxycholate, pH7.5) containing 40U/ml RNAse inhibitor (Sigma-Aldrich, St. Louis, MO) and 1X Halt™ Protease Inhibitor Cocktail (Thermofisher Scientific, Grand Island, NY). After 4-6 rounds of 50% power output sonication, 300 μl of whole cell extracts (around 500 μg total proteins) were incubated with normal rabbit IgG, Smad3, Smad4, or PPM1A antibodies (1 μg) at 4°C overnight. Next day, the immune complexes were captured with 50 μl protein A/G agarose beads (Santa Cruz Biotechnology, Dallas, TX, USA). After washing with FA lysis buffer, samples were incubated with Proteinase K at 42°C for 1 hour to digest the protein and then immunoprecipitated RNA was isolated. Purified RNA was subjected to qRT-PCR analysis for detecting the presence of GAS5.

### Western blot

Cultured cells or tissue samples were lysed in RIPA buffer (50 mM Tris-HCl, pH 7.4; 150 mM NaCl; 1% NP-40; and 0.1% SDS), and incubated with continuous rotation for 10 min at 4 °C, and then centrifuged at 12 000 ×g. The supernatant was collected, and the protein concentration was determined by a BCA assay (Pierce, Rockford, USA). Protein extracts (60-100 µg) were dissolved on 10% sodium dodecyl sulfate-polyacrylamide gels (SDS-PAGE) and transferred to polyvinylidene difluoride (PVDF) membranes. The membranes were blocked with 5% non-fat milk in Tris-buffered saline (TBS) plus Tween-20 (TBST) at room temperature for 1 h followed by incubation with primary antibodies diluted in TBST at 4 °C overnight. After three 10-min washing with TBST, blots were incubated with the appropriate secondary antibody conjugated to HRP at room temperature for 1 h. The protein expression was detected with enhanced chemiluminescent reagent.

### Co-immunoprecipitation assay (Co-IP)

Cells were transduced with Ad-GFP or Ad-GAS5 for 24 hours, and lysed with ice-cold lysis buffer containing protease inhibitor mix (Sigma). The lysates were incubated with IgG or anti-PPM1A antibody for 1 h followed by incubation with protein A/G-beads at 4 °C for 12 h. The immunoprecipitates were pelleted, washed, and subjected to immunoblotting.

### Chromatin immunoprecipitation (ChIP)

Fresh tissues were minced into small pieces in ice-cold PBS with a clean razor blade. Formaldehyde was then added to a final concentration of 1% and incubated with shaking at room temperature for 10 min. The cross-linked tissues were collected by centrifugation at 4°C and washed with PBS containing protease inhibitors before final collection. The tissues were resuspended by rotating in 1% SDS lysis buffer at 4°C for 20 min followed by sonication on ice to shear DNA into 500-1000 bp fragments. The lysates were immunoprecipitated with 2 μg of IgG (negative control) or Smad3 antibody in coimmunoprecipitation reagents (17-195, Millipore). Semi-quantitative and quantitative PCR were performed to amplify α-SMA or Col1A promoter regions containing SBE.

### Luciferase reporter assay

3T3 cells cultured in 12-well plates were transduced with AdGFP or AdGAS5 and transfected with 250 ng of firefly luciferase reporter plasmid driven by α-SMA promoter using Lipofectamie LTX (Invitrogen, USA). Cells were treated with vehicle or 5 ng/ml of TGF-β1 for 8 hours, and luciferase activities were measured using a luciferase assay kit (Promega) by following the manufacturer’s protocol. The experiments were repeated for three times with triplicates.

### Immunohistochemistry (IHC) staining

Tissue sections were rehydrated, permeabilized with 0.01% Triton X-100 in PBS, blocked with 10% goat serum, and incubated with primary antibodies at 4°C overnight followed by incubation with horseradish peroxidase (HRP)-conjugated secondary antibody. The sections were counterstained with hematoxylin.

### MTT cell proliferation assay

Cell proliferation was evaluated with 3-(4,5-dimethylthiazol-2-yl)-2, 5-diphenyltetrazolium (MTT) assay using a TACS MTT Cell Proliferation Assay Kit (Trivegen). The optical density at 570 nm was measured.

### Fluorescence in situ hybridization (FISH)

GAS5 RNA probes (sense/antisense) were synthesized and labeled using the FISH Tag RNA Multicolor kit (Life Technologies, Gaithersburg, MD). Skin tissue cryosections were digested with 20 µg/ml of proteinase K at 37 °C for 1 hour, and washed with 2 × SSC solution and with water for 5 min each at room temperature. The slides were dehydrated, air-dried and incubated with pre-denatured GAS5 probes in a dark and humid environment for hybridization at 55°C for 24 hours. The slides were then washed in 50% formamide in 2x SSC for 4 times before mounting. Nuclear was counterstained with 5,6-diamidino-2-phenylindole (DAPI).

### Statistical analysis

Sample or experiment sizes were determined empirically to achieve sufficient statistical power. For animal studies, the sample size was chosen to minimize the number of sacrificed animals while obtaining sufficient statistical power. In all of the experiments reported in this study, no data point was excluded. No randomization was used in this study. There was no blinding method used to assign individuals to experimental groups. The variance is similar between groups that are being statistically compared.

All in vitro experiments is repeated at least three times in triplicates. At least five mice were used for each treatment group in bleomycin-induced skin fibrosis studies. All values are presented as means ± SEM. Comparisons of parameters between two groups were made by two-tailed Student’s t-tests. The differences among several groups will be evaluated by one-way ANOVA with Tukey-Kramer post hoc evaluation. p-values <0.05 will be considered statistically significant. P values less than 0.05 and 0.01 were considered significant (*) or very significant (**), respectively.

## Results

### GAS5 expression was decreased in TGF-β-activated fibroblasts

TGF-β activated fibroblasts as shown by the induction of myofibroblast marker α-SMA and Col1A in both 3T3 fibroblasts and primary cultured mouse skin fibroblasts (Fig. 1A-1D). To test if GAS5 is involved in the fibroblast activation, we first examined GAS5 expression in TGF-β-treated fibroblasts. 3T3 fibroblasts were treated with vehicle, 1, 2, 5, or 10 ng/ml TGF-β for 12 hours in serum-free DMEM. Total RNA was extracted, and GAS5 expression was detected by RT-qPCR. As shown in Fig. 1E, GAS5 was down-regulated in TGF-β-treated 3T3 cells. Interestingly, 10 ng/ml of TGF-β slightly increased the GAS5 expression as compared to the treatment with 5 ng/ml of TGF-β (Fig 1E). This is probably because high concentration of TGF-β (≥10 ng/ml) can inhibit cell proliferation, which could re-activate GAS5 transcription as a feedback response. TGF-β-induced GAS5 reduction was also validated in primary cultured mouse skin fibroblasts (Fig. 1F). Since fibroblast activation is an essential process leading to tissue fibrosis, and bleomycin-induced skin fibrosis involves TGF-β signaling (13, 19, 43), we assessed if GAS5 expression is altered in bleomycin-treated skin tissues. C57BL/6 mice were injected subcutaneously with 0.02 U bleomycin every other day for 14 days, skin tissues were dissected, tissue RNA was extracted, and GAS5 expression was detected by qPCR. As shown in Fig. 1G, GAS5 was indeed downregulated in fibrotic skin tissues. Moreover, we observed GAS5-expressing cells by fluorescence in situ hybridization (FISH) assay. Comparing to the vehicle-treated skins, the numbers of GAS5-positive cells were significantly reduced in bleomycin-treated skin tissues (Fig. 1H-1I). These results suggested that GAS5 may be involved in TGF-β-induced fibroblast activation and skin fibrosis.

**Figure 1:**
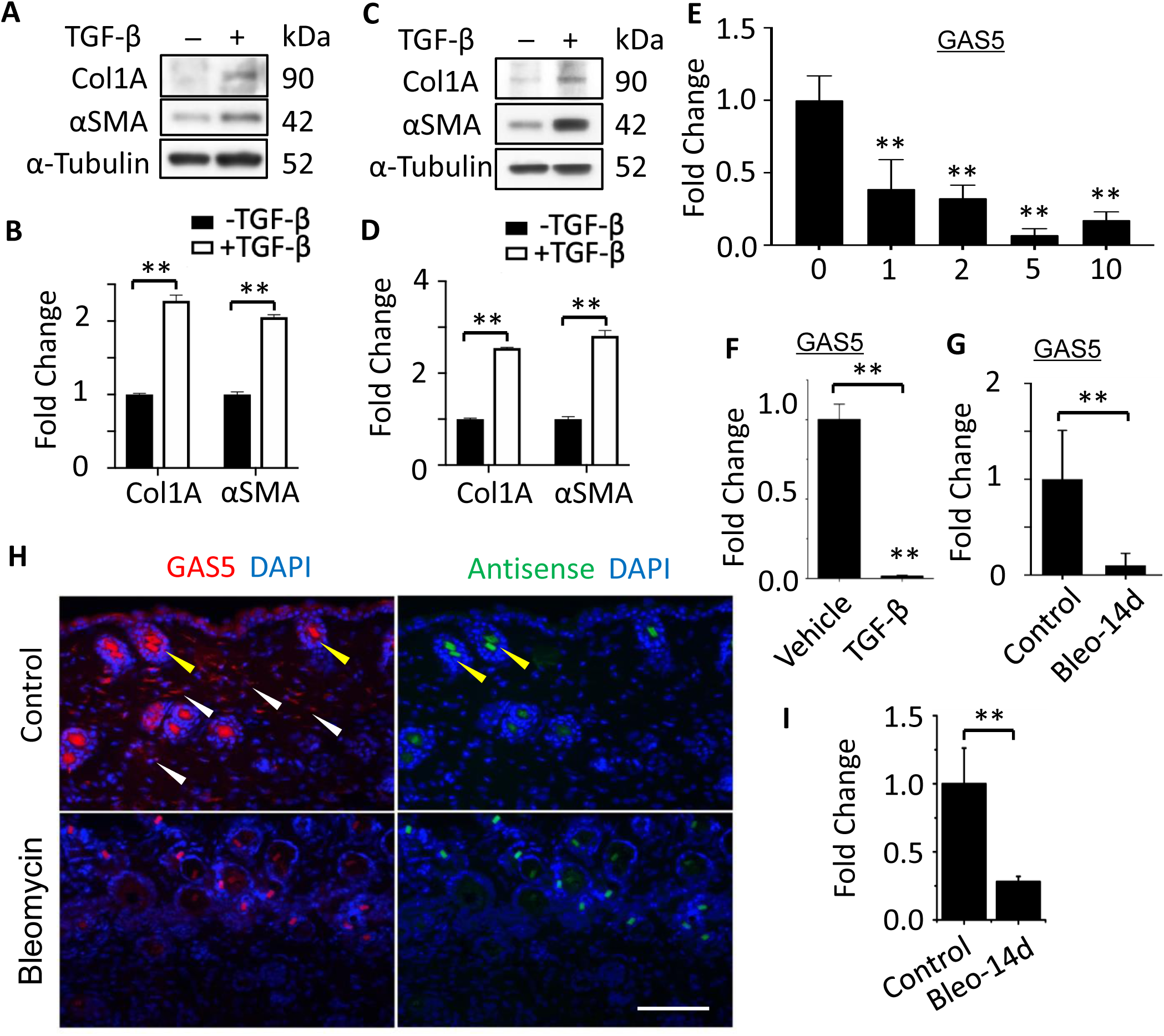
LncRNA GAS5 was downregulated in fibroblasts by TGF-β. **A-D)**, TGF-β (5ng/ml) induced the expression of myofibroblast cell marker Col1A and αSMA in 3T3 fibroblasts (A-B) and primary skin fibroblasts (C-D). **B)** Quantification of protein expression in **A** by normalizing to α-Tubulin. **D)** Quantification of protein expression in **C** by normalizing to α-Tubulin. **E)** TGF-β down-regulated GAS5 in 3T3 fibroblasts in a dose-dependent manner. 3T3 fibroblasts were cultured in serum-free DMEM with 0, 1, 2, 5, or 10 ng/ml of TGF-β for 12 hrs. GAS5 was detected by RT-qPCR. **F)** TGF-β (5ng/ml) down-regulated GAS5 in primary mouse skin fibroblasts. **G)** GAS5 was down-regulated in skin tissues with bleomycin treatment, as analyzed by RT-qPCR. C57BL/6 mice were injected subcutaneously with bleomycin (Bleo; 0.02 U) every other day for 14 days. GAS5 RNA expression was calculated by normalizing to cyclophilin mRNA. **H)** GAS5 positive cells in skin tissues were decreased by bleomycin treatment. Skin tissues from (G) was fixed in 4% PFA, and GAS5 positive cells were shown by RNA-FISH staining. Positive and false-positive staining were indicated by white and yellow arrows, respectively. **I)** Quantification of GAS5-expressing cells by counting the positive staining from 10 different fields, shown as the fold change. Bar: 200μm. * p<0.05; ** p<0.01; n=3-5. All values are presented as means ± SEM. T-tests were performed for B, D, F, G, I, and one-way ANOVA test was performed for E.

### GAS5 blocked TGF-β-induced fibroblast-myofibroblast transition

Since TGF-β induces fibroblast activation through Smad-dependent pathway, and our previous studies have shown that GAS5 blocks TGF-β/Smad3 signaling in SMC differentiation, we sought to determine if GAS5 affects TGF-β-induced fibroblast to myofibroblast transition. Thus, we detected α-SMA and Col1A protein expression in 3T3 cells transduced with AdGFP or AdGAS5 along with transfection with control or GAS5 siRNA. GAS5 overexpression and knockdown efficacies were detected by RT-qPCR (Supplemental Fig. S1). As shown in Fig. 2A-2B, overexpression of GAS5 suppressed Col1A and αSMA protein expression both at the basal state and under TGF-β treatment. Conversely, knockdown of GAS5 by its siRNA increased TGF-β-induced Col1A and α-SMA expression (Fig. 2C-2D). Consistent with the protein expression, GAS5 negatively regulated Col1A and α-SMA mRNA expression (Fig. 2E-2F). Interestingly, GAS5 caused 5 times more reduction in Col1A and 7 times more reduction in α-SMA expression in TGF-β-treated cells than the vehicle-treated cells (Fig. 2E), suggesting that GAS5 regulates Col1A and α-SMA expression primarily in association with TGF-β signaling. Moreover, overexpression of GAS5 also suppressed TGF-β-induced α-SMA promoter activity (Fig. 2G). These results indicated that GAS5 inhibits TGF-β-induced fibroblast activation by negatively regulating myofibroblast marker gene transcription.

**Figure 2.**
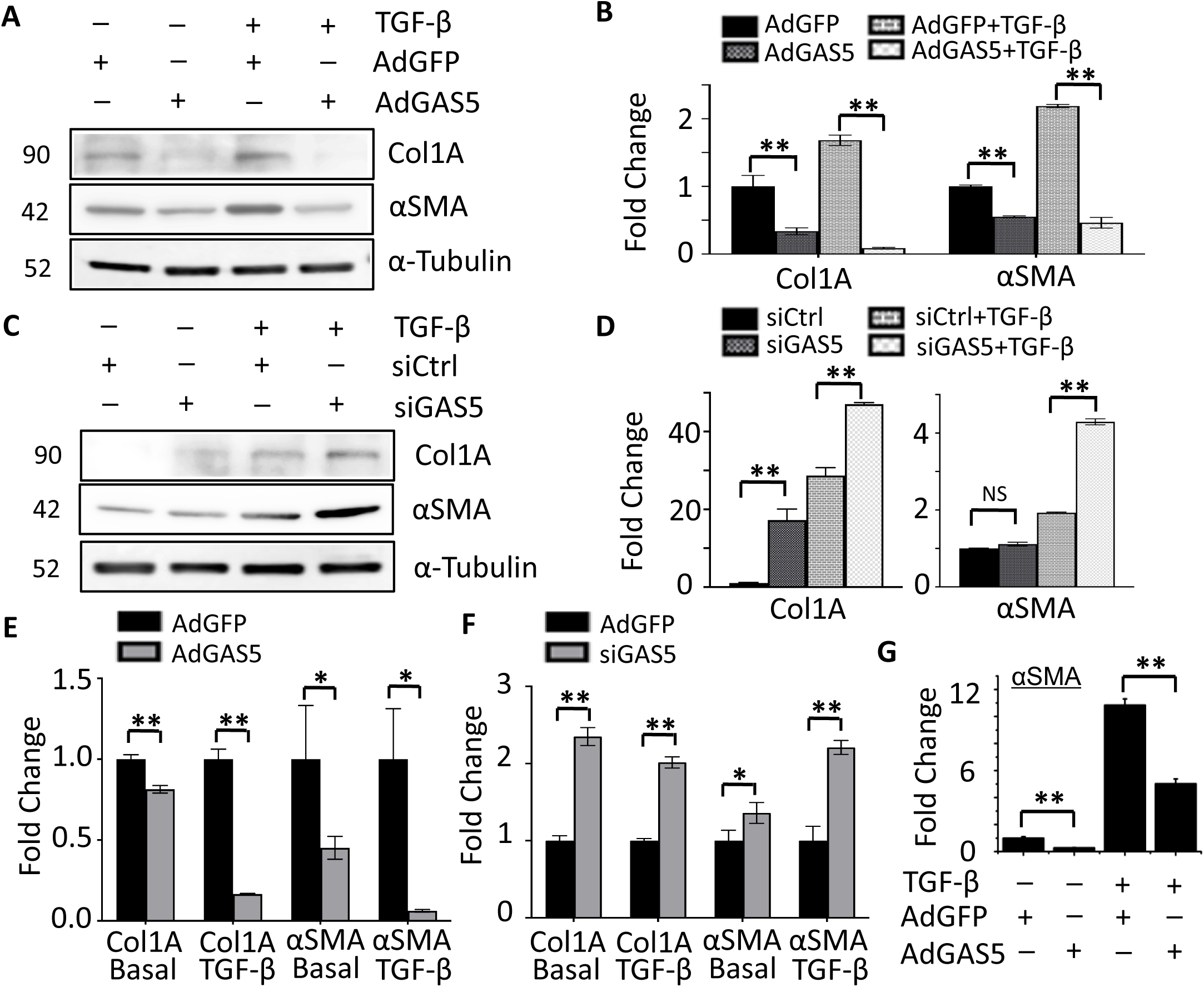
GAS5 suppressed TGF-β-induced fibroblast-myofibroblast transition. **A-B)** GAS5 suppressed αSMA and Col1A protein expression. 3T3 cells were transduced with AdGFP or AdGAS5 followed by vehicle or 5 ng/ml of TGF-β treatment for 24 hrs. αSMA and Col1A expression was assessed by Western blot (A) and quantified by normalizing to α-Tubulin (B). **C-D)** Knockdown of GAS5 increased αSMA and Col1A protein expression. 3T3 cells were transfected with siCtrl or siGAS5 followed by vehicle or 5 ng/ml of TGF-β treatment for 24 hrs. αSMA and Col1A expression was assessed by Western blot (C) and quantified by normalizing to α-Tubulin (D). **E)** GAS5 suppressed TGF-β-induced αSMA and Col1A mRNA expression. 3T3 cells were transduced with AdGFP or AdGAS5 followed by vehicle (Basal) or 5 ng/ml of TGF-β treatment for 12 hrs. mRNA expression was assessed by RT-qPCR and normalized to cyclophilin. **F)** Knockdown of GAS5 increased αSMA and Col1A mRNA expression. 3T3 cells were transfected with siCtrl or siGAS5 followed by vehicle (Basal) or 5 ng/ml of TGF-β treatment for 12 hrs. mRNA expression was assessed by qPCR and normalized to cyclophilin. **G)** GAS5 suppressed TGF-β-induced αSMA promoter activity. 3T3 cells were transduced with AdGFP or AdGAS5 and transfected with αSMA promoter luciferase reporter for 24 hours prior to the treatment with vehicle (-) or 5 ng/ml of TGF-β for 8 hours. Luciferase assays were performed. NS: not significant; * p<0.05; ** p<0.01; n=3. All values are presented as means ± SEM. one-way ANOVA tests were performed.

### GAS5 promoted Smad3 dephosphorylation

Previous studies have shown that TGF-β/Smad3 signaling is continuously activated or phosphorylated during tissue fibrogenesis, which causes sustainable Smad nuclear retention (46). We have reported that Smad3 is phosphorylated and translocated into nuclei during the initial stage of TGF-β stimulation while shuttling back to cytoplasm at the later stage of TGF-β-induced smooth muscle differentiation (55). These observations prompted us to hypothesize that GAS5 may alter Smad phosphorylation status/nuclear retention in order to regulate TGF-β-induced fibroblast activation. Since Smad nuclear localization depends on its phosphorylation status, we tested if GAS5 affects Smad2/3 phosphorylation/dephosphorylation turnover in 3T3 cells because Smad2/3 are the two major Smad proteins downstream of TGF-β signaling. As shown in Fig. 3A-3B, TGF-β induced both Smad2 and Smad3 phosphorylation. However, overexpression of GAS5 decreased Smad3, but not Smad2, phosphorylation. These data suggest that GAS5 regulates myofibroblast transition through promoting Smad3 dephosphorylation in 3T3 cells.

**Figure 3.**
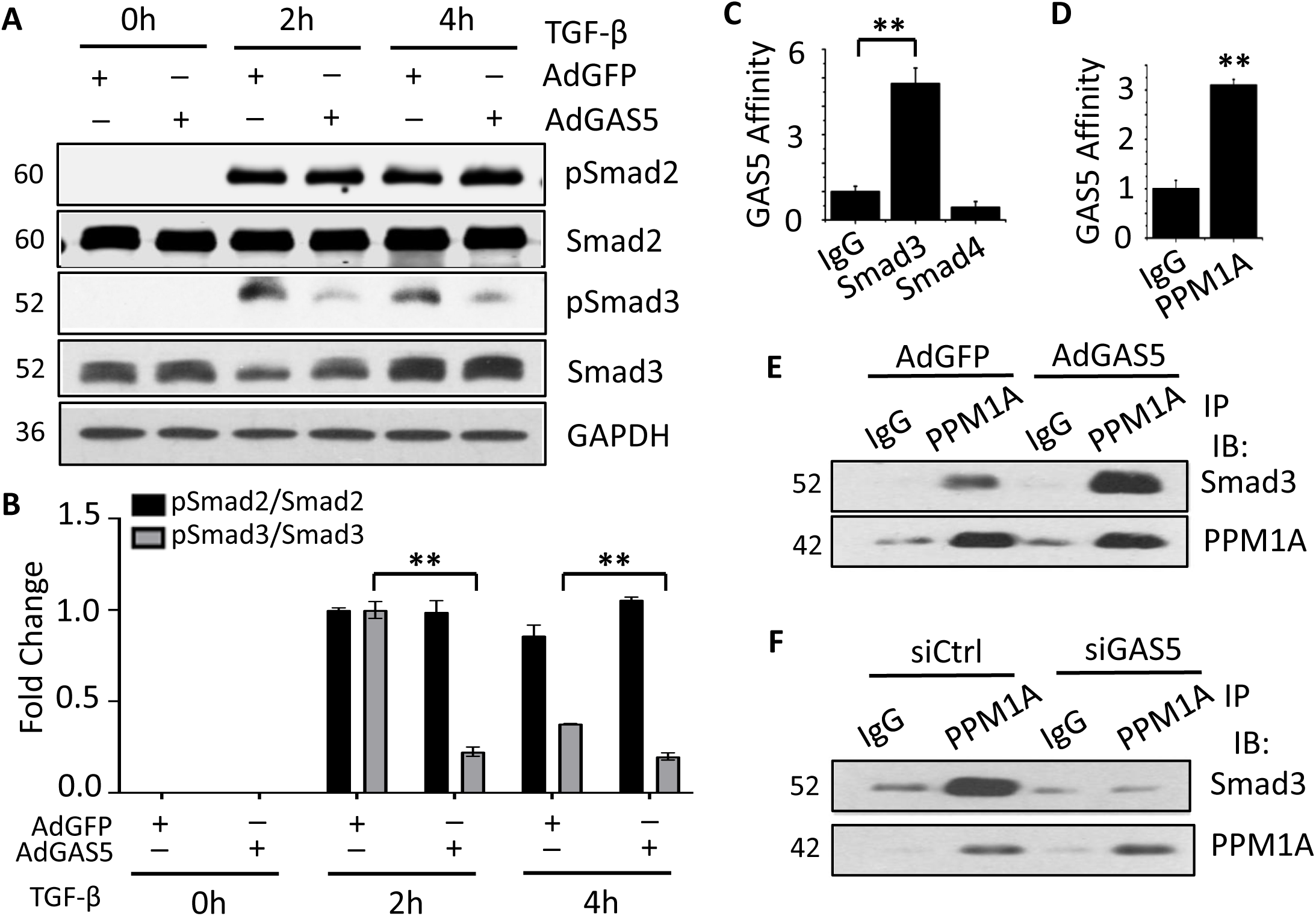
GAS5 promoted Smad3 dephosphorylation through facilitating PPM1A binding to Smad3. **A)** GAS5 accelerated Smad3 dephosphorylation in TGF-β-treated fibroblasts. 3T3 cells were transduced with AdGFP or AdGAS5 followed by vehicle (0 h) or 5 ng/ml of TGF-β treatment for 2 or 4 h. Phospho- and total Smad2 and Smad3 were detected by Western blot. **B)** Phospho-Smad2 and Phospho-Smad3 were quantified by normalizing to their total protein, respectively. **C)** GAS5 bound to Smad3 as shown by RIP assay. Smad3- and Smad4-interacting molecules in 3T3 cells were pulled down by their antibodies, respectively. The presence of GAS5 was detected via qPCR. **D)** GAS5 bound PPM1A in 3T3 fibroblasts. PPM1A-interacting molecules in 3T3 cells was pulled down by its antibody, and the presence of GAS5 was detected by qPCR. **E-F)** Overexpression of GAS5 promoted while Knockdown of GAS5 suppressed Smad3-PPM1A interaction. 3T3 cells were transduced with AdGFP or AdGAS5 (E) or transfected with siCtrl or siGAS5 (F) as indicated for 24 h. Co-immunoprecipitation was preformed by using PPM1A antibody, and the presence of Smad3 was detected by Western blot. ** p<0.01; n=3. All values are presented as means ± SEM. One-way ANOVA tests were performed for B, C, and T-test was performed for D.

### GAS5 bound to Smad3 to increase Smad3 binding to PPM1A

Smad phosphorylation status in the nuclei is controlled by Smad phosphatase (8, 10, 15). To determine the mechanism by which GAS5 promotes Smad dephosphorylation, we first predicted the interactions between GAS5 and various known Smad phosphatases through lncPro (http://bioinfo.bjmu.edu.cn/lncpro/) (31). As shown in supplementary Table 2, PPM1A is the top candidate binding GAS5. Since GAS5 only binds Smad3, but not Smad2 (49), and GAS5 overexpression caused Smad3, but not Smad2, dephosphorylation (Fig 3A-3B), we sought to test how GAS5 affects Smad3 phosphorylation and hypothesized that GAS5 regulates Smad3 turnover through binding both Smad3 and PPM1A. Thus, we experimentally determined if GAS5 interacts with PPM1A and Smad3 by performing RNA immunoprecipitation (RIP) assays. RNA-protein complex from 3T3 cells were pulled down using anti-Smad3 or anti-PPM1A antibody with IgG used as a negative control. The presence of GAS5 in the complex was detected by RT-qPCR. As shown in Fig. 3C-3D, GAS5 directly bound to Smad3 (Fig. 3C) as well as PPM1A (Fig. 3D). To test if Smad3 interacts with PPM1A in 3T3 fibroblasts and determine if GAS5 affects Smad3-PPM1A interaction, co-immunoprecipitation (Co-IP) was performed using anti-PPM1A antibody and cell lysates isolated from 3T3 cells transduced with AdGFP/AdGAS5 or transfected with siCtrl/siGAS5. As shown in Fig. 3E-3F, Smad3 indeed interacted with PPM1A. And importantly, overexpression of GAS5 enhanced Smad3-PPM1A interaction while knockdown of GAS5 suppressed Smad3-PPM1A interaction. These results indicated that GAS5 causes Smad3 dephosphorylation by promoting PPM1A binding to Smad3.

### GAS5 inhibited myofibroblast activation through PPM1A-mediated Smad3 dephosphorylation

To determine if PPM1A is important for GAS5 function in Smad3 signaling during myofibroblast activation, we knocked down PPM1A in 3T3 cells by its siRNA along with GAS5 overexpression and TGF-β treatment. As shown in Fig. 4A-4B, overexpression of GAS5 down-regulated Col1A and α-SMA expression as well as Smad3 phosphorylation. However, knockdown of PPM1A rescued, at least partially, the inhibitory effect of GAS5 on Smad3 phosphorylation and Col1A and α-SMA expression, indicating that PPM1A mediated GAS5 function in Smad3 phosphorylation during the myofibroblast activation.

**Figure 4.**
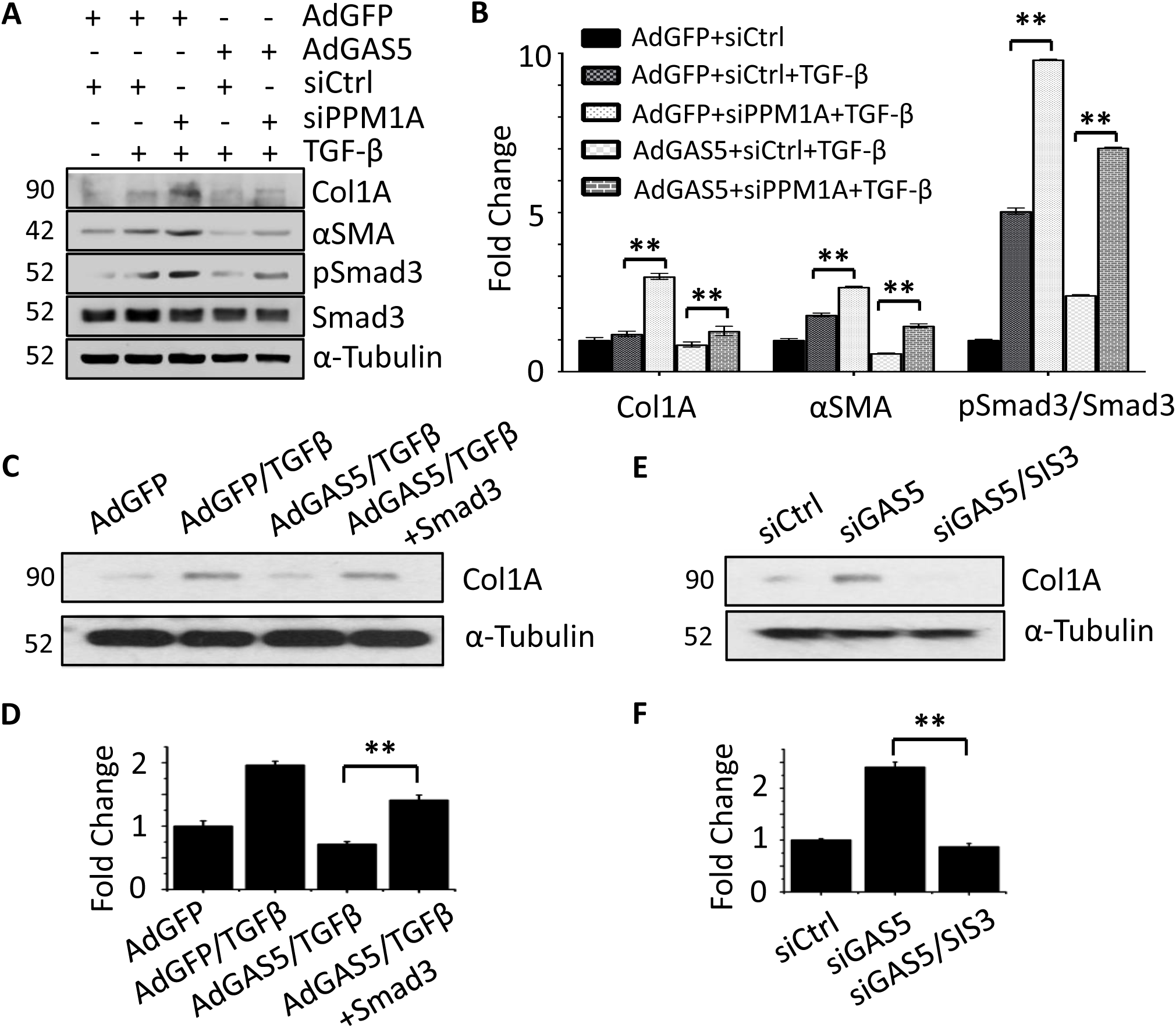
GAS5 regulated myofibroblast transition through PPM1A-induced Smad3 dephosphorylation. **A-B)** Knockdown of PPM1A rescued the inhibitory effect of GAS5 on Smad3 phosphorylation and myofibroblast marker gene expression. 3T3 cells were transduced with AdGFP or AdGAS5 along with transfection of siCtrl or siPPM1A followed by vehicle (-) or TGF-β treatment (5 ng/ml) for 24 hours. Protein expression was assessed by Western blot (A) and quantified by normalizing to α-Tubulin (for αSMA and Col1A) or total Smad3 (for pSmad3) (B). **C-D)** Overexpression of Smad3 rescued GAS5-blocked Col1A expression. 3T3 cells were transduced with AdGFP or AdGAS5 and transfected with control or Smad3 expression plasmid followed by vehicle or TGF-βtreatment (5 ng/ml) for 24 hours. Col1A expression was assessed by Western blot (C) and quantified by normalizing to α-Tubulin (D). **E-F)** Smad3 inhibitor SIS3 blunted GAS5 knockdown-enhanced Col1A expression. 3T3 cells were transfected with siCtrl or siGAS5 followed by vehicle or SIS3 (10 μM) treatment for 24 hours. Col1A expression was assessed by Western blot (E) and quantified by normalizing to α-Tubulin (F). ** p<0.01; n=3. All values are presented as means ± SEM. One-way ANOVA tests were performed.

To determine if GAS5 regulates myofibroblast gene expression through Smad3, we overexpressed GAS5 in 3T3 cells via adenoviral transduction along with transfection of Smad3 expression plasmid followed by TGF-β induction. As shown in Fig. 4C-4D, overexpression of GAS5 suppressed TGF-β-induced Col1A expression in 3T3 cells. However, Smad3 overexpression restored the Col1A expression. On the other hand, knockdown of GAS5 increased Col1A expression, but blockade of Smad3 activity by its inhibitor SIS3 demolished the effect of silencing GAS5 on Col1A expression (Fig. 4E-4F). These results demonstrated that GAS5 negatively regulated the myofibroblast gene expression by suppressing PPM1A-induced Smad3 dephosphrylation.

### GAS5 suppressed TGF-β induced fibroblast proliferation

Fibroblast proliferation is one of the main sources for the tissue mass increase during fibrosis (12, 18). TGF-β signaling is known to regulate cell proliferation (1, 4, 24). To confirm the function of TGF-β in fibroblast proliferation, 3T3 cells were treated with 0, 1, 5 or 10 ng/ml TGF-β for 48 hours in serum-free DMEM, following by MTT assay to assess cell proliferation. As shown in Supplemental Fig. S2A, TGF-β increased 3T3 cell proliferation in a dose-dependent manner. Consistently, PCNA expression was also dose-dependently increased in response to TGF-β (Fig. S2B-2C). To determine the role of GAS5 in TGF-β-induced 3T3 proliferation, we overexpressed GAS5 by transducing 3T3 cells with control or adenoviral vector expressing GAS5 (AdGAS5) followed by treatment with vehicle or 5 ng/ml of TGF-β for 48 hours for MTT assay or 24 hours for Western blot. As shown in Figure S2D-2F, overexpression of GAS5 suppressed TGF-β-induced 3T3 proliferation (Fig. S2D) as well as PCNA protein expression (Fig. S2E-2F). On the other hand, knockdown of GAS5 by its siRNA increased 3T3 cell proliferation (Fig. S2G) and increased PCNA protein expression (Fig. S2H-2I). These results indicated that GAS5 suppressed TGF-β-induced fibroblast proliferation.

### GAS5 suppressed fibroblast proliferation through TGF-β/JNK signaling

It is known that TGF-β/Smad signaling cross-talks with multiple non-Smad signaling pathways, most often PI3K/Akt, JNK and p38 pathways, to regulate cell proliferation (2, 3, 17, 23, 41, 59). To determine if GAS5 inhibits fibroblast proliferation by targeting these signaling pathways, we assessed the phosphorylation of key components in PI3K/Akt, JNK and p38 signaling pathways in TGF-β-treated 3T3 cells. As shown in Supplemental Figure S3A-3B, TGF-β increased Akt and JNK phosphorylation, suggesting these signaling pathways were activated during TGF-β-induced fibroblast proliferation. However, comparing to AdGFP-transduced cells, overexpression of GAS5 (AdGAS5) suppressed JNK phosphorylation while had no effect on p38 phosphorylation and increased the Akt phosphorylation (Fig. S3A-3B). These results suggested that GAS5 suppressed TGF-β-induced fibroblast proliferation only through JNK pathway. To further validate the function of JNK signaling in GAS5-regulated fibroblast proliferation, we knocked down GAS5 in 3T3 cells via siRNA transfection followed by treatment with vehicle or JNK pathway inhibitor, SP600125. As shown in Fig. S3C, knockdown of GAS5 increased 3T3 cell proliferation, but JNK inhibitor reversed the effect of GAS5. These results demonstrated that GAS5 inhibits fibroblast proliferation through interfering with TGF-β-JNK pathways.

### In vivo local delivery of GAS5 via adenoviral vector suppressed skin fibrosis in mice

Since fibroblast activation is an important mechanism underlying pathological tissue/organ fibrosis, we tested the function of GAS5 in skin fibrosis by using a bleomycin-induced TGF-β-dependent skin sclerosis model (43). GAS5 was overexpressed via adenoviral delivery in bleomycin-treated mouse skin. As shown in Fig. 5A-5D, overexpression of GAS5 attenuated bleomycin-induced skin sclerosis as indicated by the reduction of both skin thickness (Fig. 5A-5B) and skin tissue collagen deposition (Fig. 5C-5D). Immunohistochemistry (IHC) staining showed that Col1A-(Fig. 5E-5F) and αSMA-positive cells (Fig. 5G-5H) were significantly reduced in AdGAS5-transduced skin tissues, suggesting that GAS5 inhibited bleomycin-caused skin fibrosis. Moreover, overexpression of GAS5 decreased Col1A and α-SMA protein expression (Fig. 6A-6B) in bleomycin-treated skins. Importantly, GAS5 reduced the Smad3 binding to the Col1A and α-SMA promoters in a chromatin setting in vivo in the bleomycin-treated skin tissues (Fig. 6C-6F), confirming that GAS5 impedes skin fibrogenesis by inhibiting Smad3-mediated transcription of myofibroblast genes.

**Figure 5.**
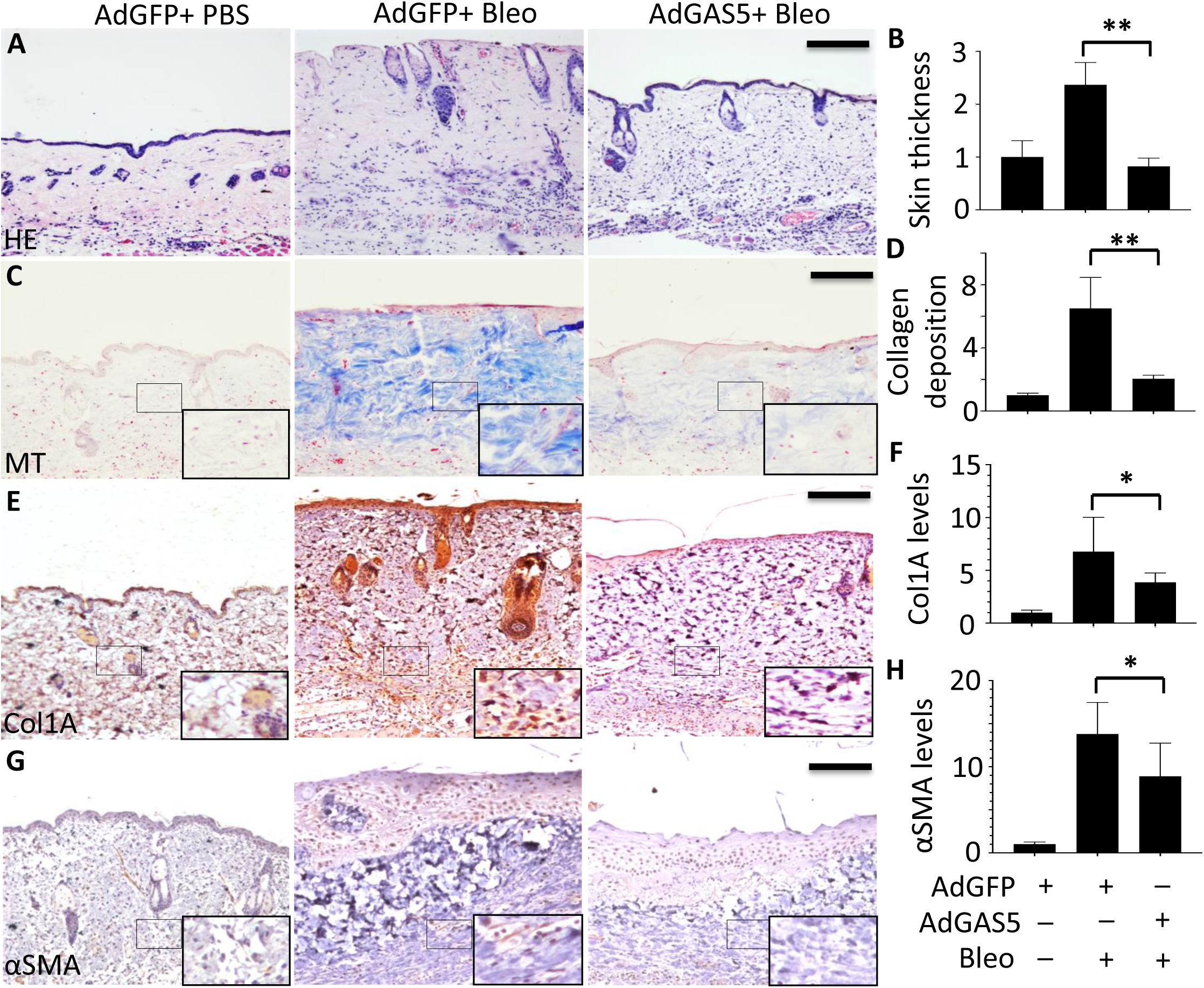
GAS5 suppressed skin fibrosis in mice. **A-D)** Forced expression of GAS5 via local adenoviral delivery suppressed bleomycin-induced skin fibrosis in mice. Mouse skins were treated with vehicle (PBS) or bleomycin (0.02U/day) every other day for 28 days. The skin tissues were stained with H&E for structural changes (A), Masson’s trichrome (MT) for collagen deposition (C). Bar: 200 μm. Skin thicknesses shown in A were averaged from 10 different fields (B), and collagen deposition in C was quantified by measuring the staining intensity from 10 different fields (D). **E-H)** Forced expression of GAS5 via local adenoviral delivery suppressed bleomycin-induced expression of Col1A (E-F) and αSMA (G-H). Bar: 100 μm. Skin tissue sections underwent immunohistochemistry (IHC) staining using Col1A (E) and αSMA (G) antibodies, respectively. Col1A (F) and αSMA (H)-positive cells were averaged from 10 different fields. The large rectangle inserts are enlarged images from the small rectangle boxes in C, E and G, respectively. * p<0.05; ** p<0.01; n=5. All values are presented as means ± SEM. One-way ANOVA tests were performed.

**Figure 6.**
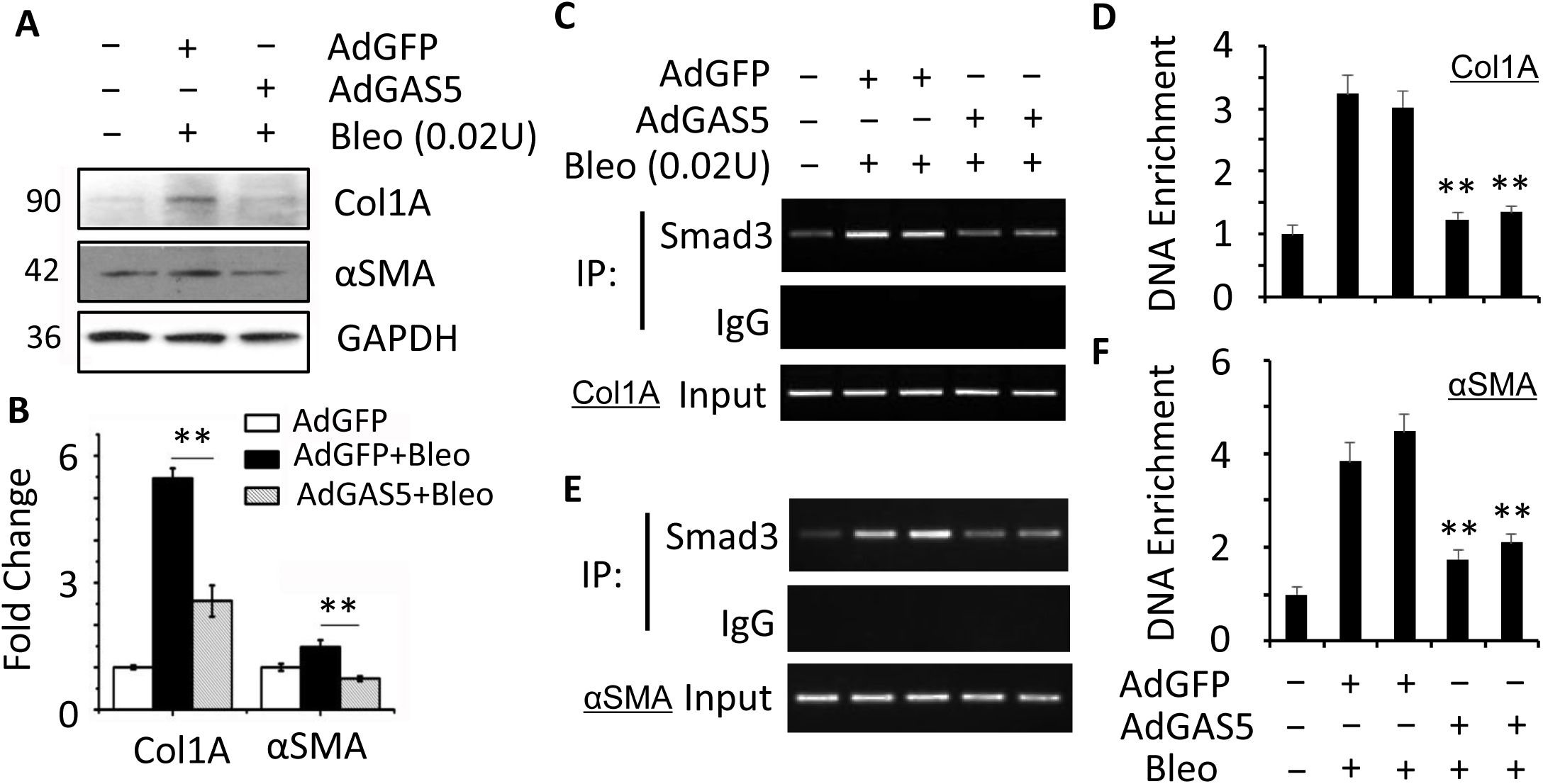
GAS5 inhibited Col1A and αSMA protein expression in fibrotic skin tissues and attenuated Smad3 binding to their promoters in vivo. **A)** GAS5 suppressed Col1A and αSMA protein expression in skin tissues with bleomycin-induced fibrosis. **B)** Col1a and αSMA protein levels in (A) were quantified by normalizing to GAPDH. ** p<0.01; n=5. **C-F)** GAS5 suppressed Smad3 binding to Col1a (C-D) and α-SMA (E-F) promoters in vivo that were significantly enriched during bleomycin-induced skin fibrosis. Smad3 binding to Col1a (C) and α-SMA (E) promoters in a chromatin setting was measured by in vivo chromatin immunoprecipitation (CHIP) assay. Smad3 binding enrichments were quantified by qPCR relative to the Smad3 binding in vehicle-treated skin tissues (D-F). ** p<0.01 vs AdGFP-treated groups; n=5. All values are presented as means ± SEM. One-way ANOVA tests were performed.

## Discussion

Fibrosis is a chronic wound healing progress characterized by fibroblast activation involving fibroblast proliferation and fibroblast-myofibroblast transition (20, 42). In this study, we identified lncRNA GAS5 as an essential regulator for TGF-β-induced fibroblast activation and skin fibrosis. Specifically, GAS5 directly bound to both Smad3 and PPM1A, thus promoted Smad3 dephosphorylation and suppressed Smad3-induced myofibroblast activation. In addition, GAS5 inhibited 3T3 cell proliferation via blocking JNK signaling (Fig. 7). Importantly, Local adenoviral delivery of GAS5 effectively suppressed bleomycin-induced skin fibrosis in mice, suggesting that GAS5 may be used as a promising RNA-based therapeutic agent for treating fibrotic diseases.

**Figure 7.**
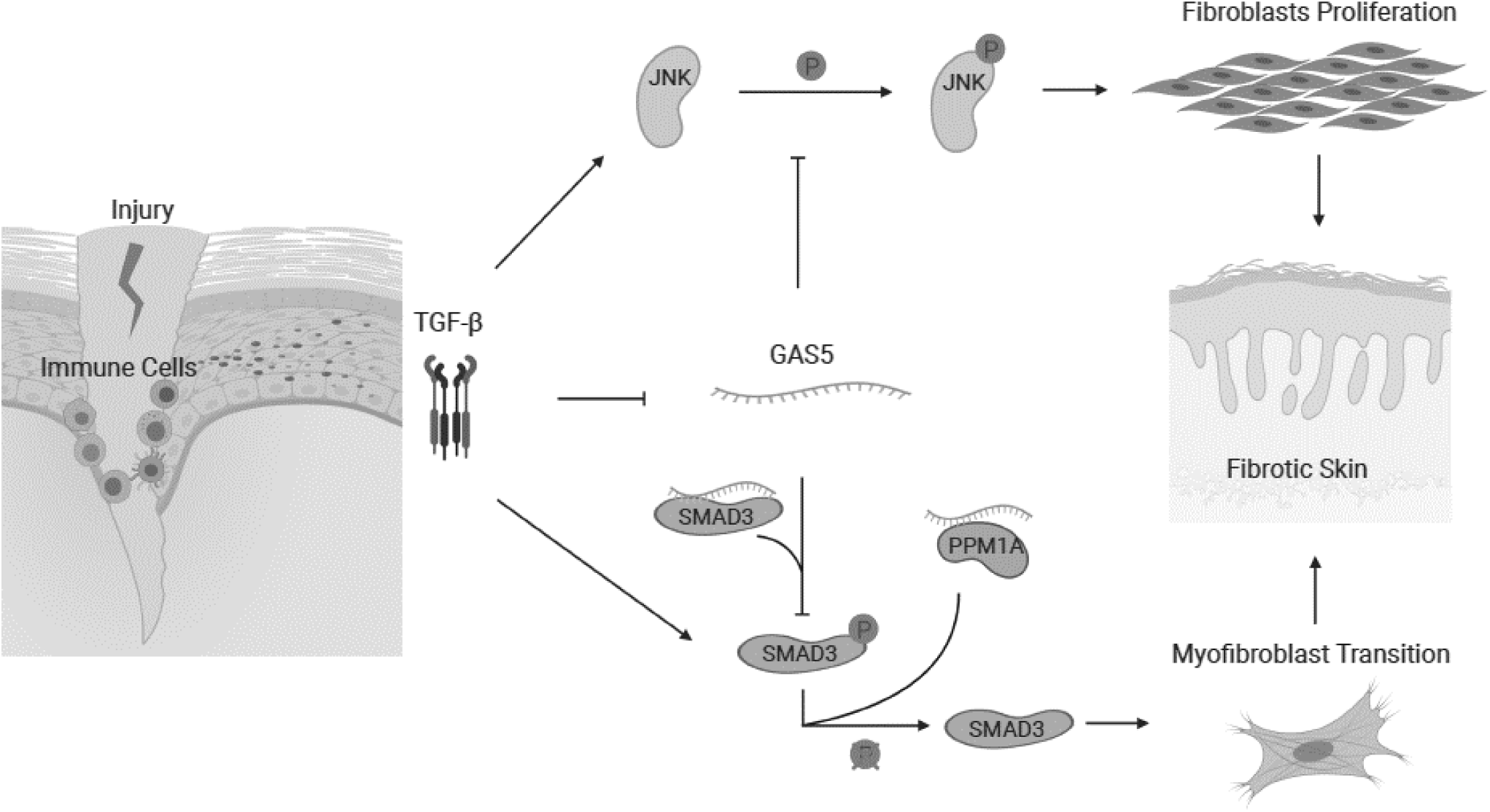
A diagram of GAS5 function in TGF-β-induced skin fibrosis. During the onset of skin fibrosis, infiltrating immune cells secret TGF-β, which activates resident fibroblasts through Smad3 signaling pathway. GAS5 suppresses the progression of fibrosis by promoting Smad3 dephosphorylation through facilitating PPM1A binding to Smad3, thus inhibiting Smad3-mediated myofibroblast activation. GAS5 also inhibits fibroblast proliferation through impeding JNK signaling pathway.

LncRNA normally functions as a suppressive protein sponge for transcription factors (27, 49). lncRNA can also serve as a scaffold RNA which brings spatial proximity of different components thus facilitates protein complex formation and enhances their functions (45, 52). GAS5 has been shown to regulate different signaling cascades through recruiting protein co-factors to form RNA-Protein complex (RNP). Indeed, we have previously identified GAS5 as a Smad3 sponge RNA in the initial stage of TGF-β signaling transduction. In the current study, we identified a new function of GAS5 at a later stage of TGF-β signaling. I.e., GAS5 can quench the persistent TGF-β/Smad signaling observed in fibrosis by promoting PPM1A-mediated Smad3 dephosphorylation. Although transient TGF-β activity is important for tissue repair/regeneration, persistent TGF-β signaling results in fibrosis and scarring in fibrotic disease (11, 30). Our results suggested that GAS5 may act as a brake for the persistent TGF-β signaling in tissue fibrosis. GAS5 affects myofibroblast marker gene Col1A and α-SMA expression at both basal and TGF-β-treated states, suggesting that GAS5 may be required for maintaining fibroblast homeostasis of the quiescent cells. Interestingly, the effect of silencing GAS5 on α-SMA is much less than Col1A in the basal state, which is probably because α-SMA expression is also regulated by many factors other than GAS5. Of significance, GAS5 has more perfound impacts on Col1A and α-SMA expression in TGF-β-treated cells, suggesting that GAS5 may be more important and powerful in blocking the fibroblast activation and fibrosis than maintaining quiescent cell activity. Indeed, GAS5 has also been identified as an anti-fibrotic lncRNA in cardiac and hepatic fibrosis although the mechanism is mainly through functioning as a miRNA sponge (16, 50, 56). Whether GAS5 regulates TGF-β/Smad signaling through sponging miRNAs in skin fibrosis requires further studies.

TGF-β not only initiates fibroblast-myofibroblast transition but also stimulates fibroblast proliferation, usually through cross-talking with Smad independent pathways, such as PI3K/Akt, JNK and p38 (1, 4, 24). Given the fact that GAS5 is a potent anti-proliferation lncRNA, it is expected that GAS5 also suppresses TGF-β-induced fibroblast proliferation. Although previous studies have shown that TGF-β signaling interacts with almost all major cellular signaling pathways (54), our results indicate that JNK is the major downstream signaling pathway mediating GAS5 function in TGF-β-induced fibroblast proliferation. There are reports showing that knockdown of GAS5 suppresses JNK phosphorylation in primary retinal ganglion cells and a neuroblastoma cell line (34, 60). These discrepancies in GAS5 function may be due to cell type-specific effects. How GAS5 interacts with JNK signaling, and whether the interaction between GAS5 and JNK signaling is Smad3-dependent are interesting subjects for future studies.

In the progression of fibrogenesis, intensive interplays occur among different cell types. Epithelial cells, fibroblast, smooth muscle cells and immune cells all have shown to play important roles in this chronic disease (21, 28, 39). M2 macrophages, for example, is known to play important roles not only in the initial wound healing response (anti-fibrotic) but also in the later TGF-β secretion and fibroblast activation stages (pro-fibrotic) (9). Interestingly, GAS5 has been reported to inhibit M2 macrophage polarization (44). Thus, GAS5 may have many unidentified functions in fibrosis. For example, it could coordinate or modulate TGF-β signaling among different cell types during fibrogenesis. The investigation into these topics may shed new light on the RNA-based therapy in treating TGF-β-related fibrotic diseases.

## Abbreviations

TGF-β: Transforming Growth Factor β
lncRNA: Long noncoding RNA
GAS5: Growth arrest-specific transcript 5
ECM: Extracellular matrix
Col1A: Collagen 1A
TβR: TGF-β receptor
JNK: c-Jun N-terminal kinase
FISH: Fluorescence in situ hybridization
RIP: RNA immunoprecipitation
Co-IP: Co-immunoprecipitation
Smad3: Mothers against decapentaplegic homolog 3
PPM1A: Protein phosphatase 1A
α-SMA: Smooth muscle α-actin
IHC: Immunohistochemistry
RNP: RNA-Protein complex

## Acknowledgments

This work was supported by grants from National Institutes of Health (HL123302, HL119053, HL135854, and HL147313 to S.-Y.C.) and VA merit Award (I01 BX004065-1 to K. F. S). R. T. is a receipient of Stanford University School of Medicine Dean’s Postdoctoral Fellowship and a TRDRP Postdoctoral Fellowship (27FT-0044).

## Endnotes

The Supplemental Materials are available in the following link: https://doi.org/10.6084/m9.figshare.12140880

