## Supplementary Figures 1-3 for "LncRNA GAS5 attenuates fibroblast activation through inhibiting Smad3 signaling"

### Supplemental Materials

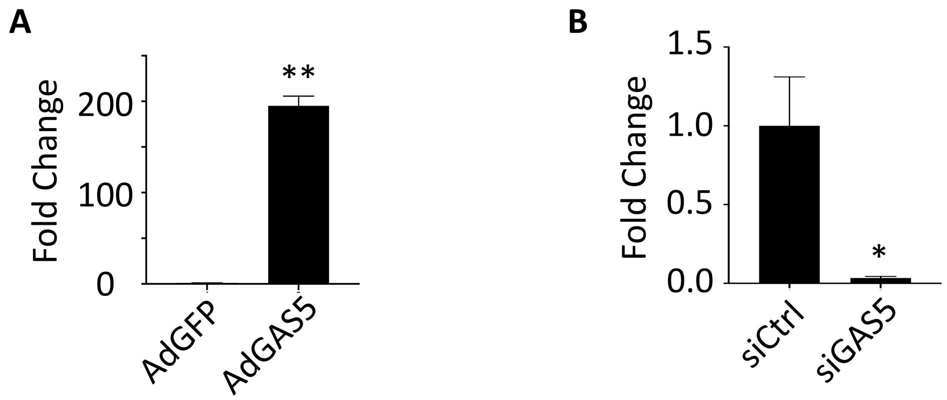


**Supplemental Figure S1. Validation of GAS5 overexpression and knockdown efficiency. A)** GAS5 adenoviral transduction (AdGAS5) dramatically increased GAS5 expression in 3T3 cells. **B)** GAS5 expression was knocked down by its siRNA (siGAS5) in 3T3 cells. *$p<0.05$; ** $p<0.01$; $n=3$.

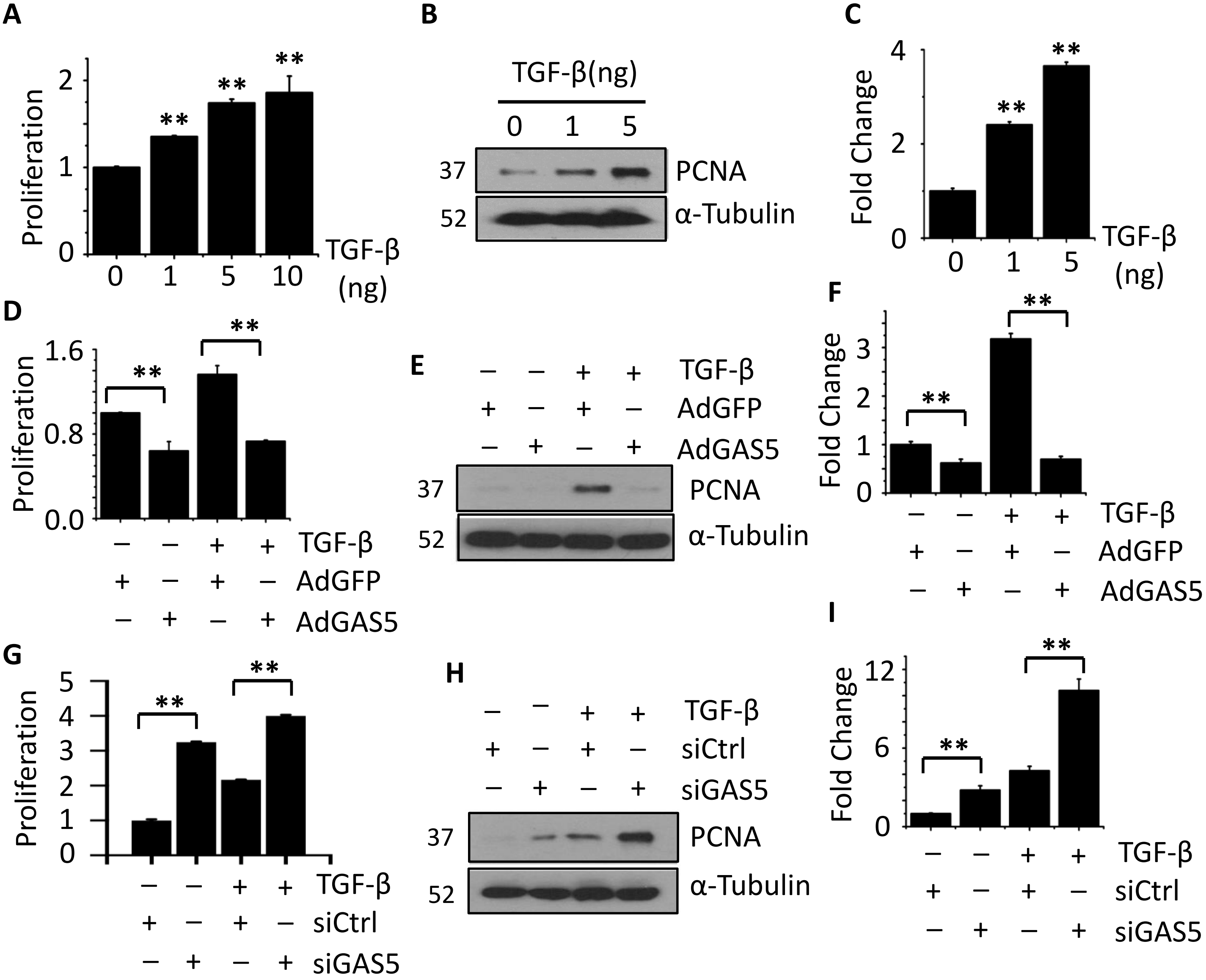


**Supplemental Figure S2. GAS5 suppressed TGF-β-induced fibroblast proliferation. A)** TGF-β induced 3T3 fibroblast proliferation in a dose-dependent manner. Cell proliferation rate was assessed by MTT assay and compared to the untreated cells (0 ng/ml). **B)** TGF-β induced PCNA expression in a dose-dependent manner in 3T3 cells. **C)** Quantification of PCNA levels by normalizing to α-Tubulin. **D-F)** Overexpression of GAS5 suppressed TGF-β-induced 3T3 cell proliferation and PCNA expression. 3T3 cells were transduced with AdGFP or AdGAS5 for 24 hrs and then cultured in DMEM with 10% FBS with vehicle or TGF-β (5 ng/ml) treatment for 48 hrs. Cell proliferation was assessed by MTT assay. PCNA expression was detected by Western blot (E) and quantified by normalizing to α-Tubulin as in (F). **G-I)** Knockdown of GAS5 promoted 3T3 proliferation and PCNA expression. 3T3 cells were transfected with siCtrl or siGAS5 for 24 hrs followed by treatment with vehicle or 5 ng/ml of TGF-β for 48 hrs. Cell proliferation and PCNA expression were assessed as in (D-F). * $p<0.05$; ** $p<0.01$; n=3.

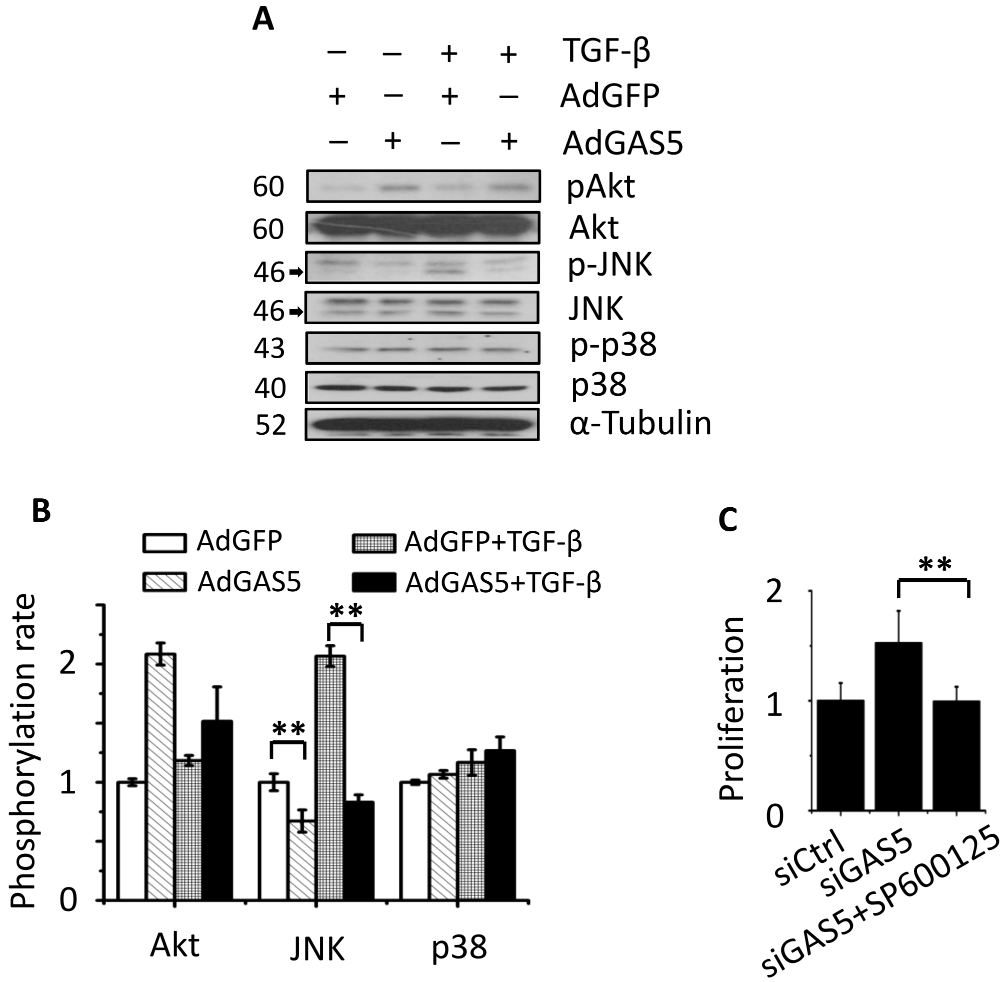


**Supplemental Figure S3.** GAS5 suppressed TGF-β- induced JNK phosphorylation. **A-B)** 3T3 cells transduced by AdGFP or AdGAS5 were treated with vehicle or 5 ng/ml of TGF-β for 12 hrs. Protein expression or phosphorylation was detected by Western blot (A). Protein phosphorylation in A was normalized to their corresponding total protein levels (B). **C)** JNK inhibitor SP600125 suppressed siGAS5-enhanced cell proliferation. siCtrl or siGAS5-transfected 3T3 cells were treated with vehicle or 10 μM JNK inhibitor SP600125 for 48 hrs. Cell proliferation was assessed by MTT assay. * $p<0.05$; ** $p<0.01$; $n=3$.

**Supplementary Table 1:** qPCR primers used in this study.

| | |
|---|---|
| mmu-GAS5-F | GGA GGT TGG TTC TGC GTG TA |
| mmu-GAS5-R | CGC ATG CTG AGT CGT CTT TG |
| mmu-SMA-F | AAT GGC TCT GGG CTC TGT AAG |
| mmu-SMA-R | CAC GAT GGA TGG GAA AAC AGC |
| mmu-Col1a-F | TCC TTC TGG TCC TCG TGG TCT C |
| mmu-Col1a-R | AGC CTC GGT GTC CCT TCA TTC C |

**Supplementary Table 2:** Smad phosphatase candidates potentially bound GAS5.

| Rank | Protein | Value |
|---|---|---|
| **1** | mmu-Ppm1a | 51.7 |
| **2** | mmu-Pdp1 | 41.5 |
| **3** | mmu-Pp2a | 37.7 |
| **4** | mmu-Mtmr4 | 37.5 |
| **5** | mmu-Scp1 | 34.9 |
| **6** | mmu-Scp2 | 27.9 |
